# Structural and functional insights into nuclear role of Parkinson’s Disease-associated α-Synuclein

**DOI:** 10.1101/2024.10.24.620072

**Authors:** Sneha Jos, Archanalakshmi Kambaru, Thazhe Kootteri Prasad, Shylaja Parthasarathi, Neelagandan Kamariah, Sangeeta Nath, Balasundaram Padmanabhan, Sivaraman Padavattan

## Abstract

α-Synuclein (αSyn) plays a critical role in the pathogenesis of ‘Synucleinopathies’. Although increased nuclear αSyn localization induces neurotoxicity, its definitive physiological role remains elusive. Previous studies on nuclear αSyn are limited to its interactions with individual histones and dsDNA, leaving a significant gap in understanding its interactions with assembled histone H2a-H2b dimer and (H3-H4)_2_ tetramer, as well as its role in chromatin regulation. Here, we demonstrated that αSyn binds specifically to both H2a-H2b and (H3-H4)_2_ with high affinity. Truncation studies revealed that αSyn(1-103) region interacts with (H3-H4)_2_, while the acidic (121-140) C-terminal end is crucial for H2a-H2b binding. Sequence analysis suggests αSyn-dimer binding region contains a conserved DEF/YxP motif present in other dimer-binding histone chaperones. High-resolution structure of αSyn- dimer binding region with H2a-H2b complex reveals that αSyn adopts two binding modes (BM1 and BM2). In BM-1, αSyn utilizes nucleosomal DNA-binding surface, while in BM-2, it engages with both DNA- and the H3-interaction interface. Additionally, dimer recognition by αSyn overlaps with other dimer-binding histone chaperones, suggesting αSyn’s potential role in the nucleosome assembly/disassembly process.

## INTRODUCTION

αSyn is a pivotal protein associated with a group of neurodegenerative diseases referred to as ‘Synucleinopathies,’ which includes Parkinson’s disease (PD), dementia with Lewy bodies (DLB), and multiple system atrophy (MSA) (Goedert, 2001). It was first identified as a neuronal protein that undergoes presynaptic and nuclear localization in electric ray fish (*Torpedo californica*) (Maroteaux *et al*, 1988). Though many cellular functions have been proposed for αSyn over the years, its precise physiological function remains unclear. In 1997, αSyn was identified as a main constituent of intracellular cytoplasmic inclusion referred to as Lewy bodies (LBs), the key pathological feature in synucleinopathies (Spillantini *et al*, 1997). Since then, most studies have focused on interconnecting αSyn aggregation properties to disease etiology (Lashuel, 2020).

Nuclear αSyn localization is associated with physio-pathology (Goers *et al*, 2003; Kontopoulos *et al*, 2006; Wakamatsu *et al*, 2007; Schell *et al*, 2009; Mbefo *et al*, 2010; Liu *et al*, 2011; Fares *et al*, 2014; Mbefo *et al*, 2015; Pinho *et al*, 2019; Davidi *et al*, 2020; Geertsma *et al*, 2022; Gonçalves & Outeiro, 2013), but less emphasis is given to understanding its specific nuclear role. Multiple lines of evidence indicate that under pathological conditions, the nuclear αSyn level increases, eliciting neurotoxicity in dopaminergic neurons and mouse models independent of its aggregation property (Wakamatsu *et al*, 2007; Goers *et al*, 2003; Kontopoulos *et al*, 2006; Geertsma *et al*, 2022). These findings raise a fundamental question regarding the mechanism of αSyn toxicity in PD: the underappreciated nuclear function versus its aggregation property. Therefore, determining αSyn’s physiological role in the nucleus is of particular interest. So far, studies on nuclear αSyn have only explored its interactions with individual core histones, linker histones, and dsDNA (Goers *et al*, 2003; Kontopoulos *et al*, 2006; Schaser *et al*, 2019; Jos *et al*, 2021). Nonetheless, how αSyn interacts with assembled histone H2a-H2b dimer, (H3-H4)_2_ tetramer, nucleosome, and its role in chromatin regulation remains unknown. In this study, we have unveiled that αSyn functions as a histone chaperone protein in the nucleus. Histone chaperones are a family of proteins that faithfully guard the histone supply chain and dynamics during replication, transcription, and DNA repair processes throughout cellular life (Warren & Shechter, 2017). Here, we investigated αSyn interaction with H2a-H2b, (H3-H4)_2_, and nucleosome core particle (NCP) using biochemical and biophysical approaches. Additionally, we determined the X-ray crystal structure of αSyn with the H2a-H2b dimer complex to 1.72 Å resolution. Remarkably, our structure revealed that the dimer recognition by αSyn overlaps with that of other chromatin regulators, suggesting a potential role in the nucleosome assembly/disassembly process. Together, these studies have provided molecular-level details and structural insights into αSyn nuclear physiological function. Based on these results, we discussed possible models for αSyn’s role in the physio-pathological conditions.

## RESULTS

αSyn belongs to the intrinsically disordered protein (IDP) family composed of 140 amino acids (14.46 kDa, pI 4.6). It consists of three domains: the positively charged amphipathic N-terminal region (1-60 residues), aggregation-prone central non-amyloid-*β* component (NAC) region (61-95 residues), and the highly acidic C-terminal tail (104-140 residues) (Stephens *et al*, 2019). The individual core histones (H2a, H2b, H3, and H4) comprise the N-terminal flexible tail and C-terminal histone-fold region and are assembled into heterodimers with complementary histones (McGinty & Tan, 2015). We have recombinantly expressed and purified human αSyn(full-length; FL), a series of C-terminal truncated αSyn constructs, and individual human core histones with/without N-terminal flexible tail as previously reported (Jos *et al*, 2021) **(Figure 1A)**. Additionally, we have purified a single-chain tailless *Xenopus laevis* H2a-H2b dimer (ScH2a-H2b) generated by linking the C-terminal end of H2b(34-126) with the N- terminal of H2a(13-102) for structural studies (Warren *et al*, 2020). It is worth noting that ScH2a-H2b dimer precipitates below 1.0 M NaCl concentration, whereas the H2a-H2b dimer and (H3-H4)_2_ tetramer assembled using individually purified core histones under denaturing conditions are soluble at physiological salt concentration (150 mM). Hence, except for structural studies, the H2a-H2b dimer and (H3-H4)_2_ tetramer used in biochemical and biophysical studies were individually purified and assembled as reported (Tanaka *et al*, 2004; Dyer *et al*, 2004).

**Figure 1:**
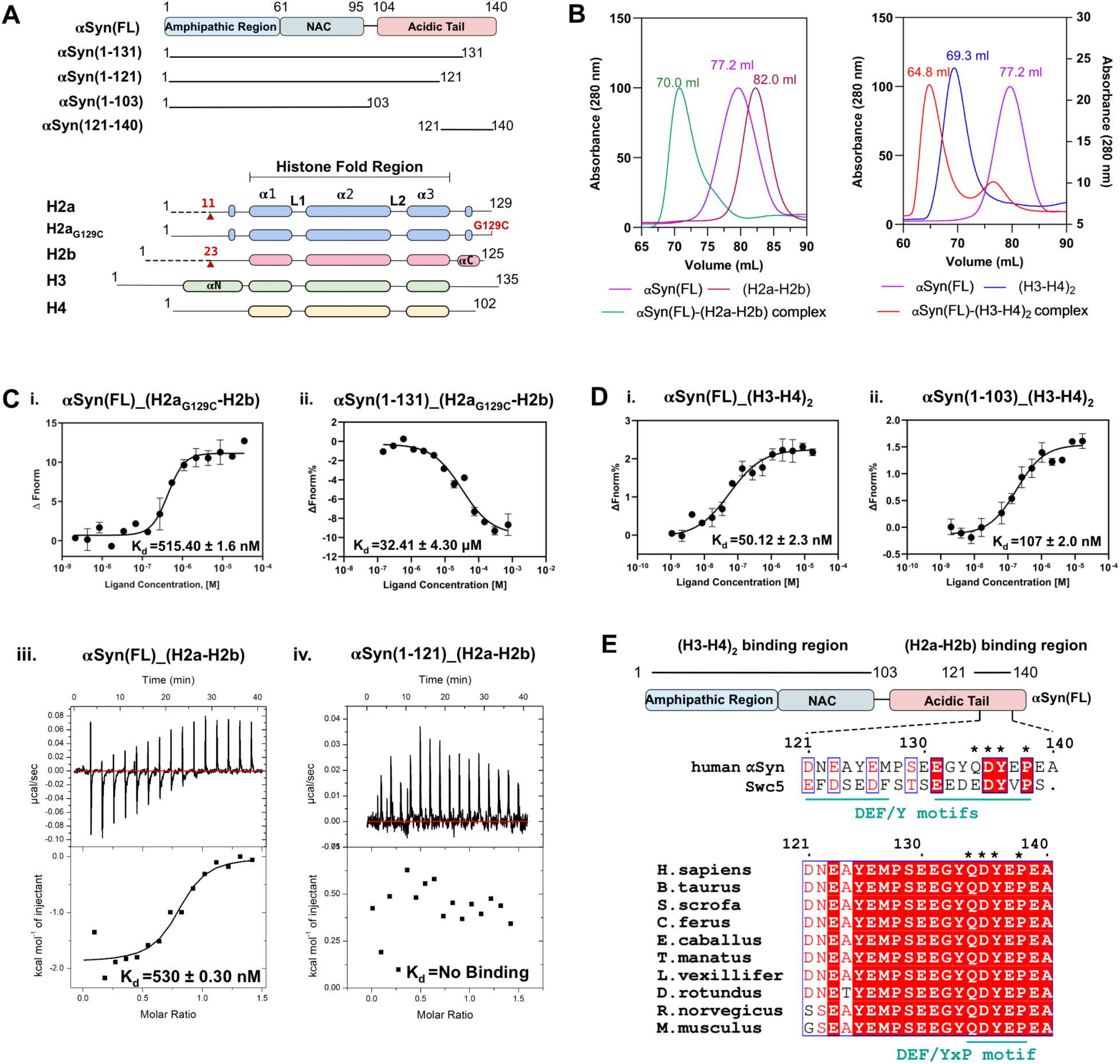
Domain analysis of αSyn binding with H2a-H2b and (H3-H4)_2_. **A.** Schematic representation of αSyn and histone constructs used in domain analysis. The red arrow indicates the H2a and H2b N-terminal tail truncation boundary. **B.** Size-exclusion chromatography shows αSyn(FL) forms a complex with (H2a-H2b) dimer (left) and (H3-H4)_2_ tetramer (right). **C.** MST analysis of αSyn(FL) (i) and αSyn(1-131) (ii) with fluorescently labeled H2a_G129C_-H2b dimer. iii. ITC analysis of the αSyn(FL) and αSyn(1-121) with H2a-H2b dimer. **D.** MST analysis of the αSyn(FL) (i) and αSyn(1-103) (ii) with fluorescently labeled (H3-H4)_2_ tetramer. For both MST and ITC, the *k_d_* ± SD values are displayed, and error bars represent SD (N=3). **E.** Sequence alignment of the human αSyn-dimer recognition region (121-140) with Swc5 in yeast (top) and multiple sequence alignment with other species (bottom). The DEF/YxP motif is indicated by (*) on top of the sequence analysis.

### αSyn forms complex with both H2a-H2b dimer and (H3-H4)_2_ tetramer

To examine whether αSyn(FL) binds assembled H2a-H2b dimer and (H3-H4)_2_ tetramer, we independently reconstituted αSyn(FL) with these assembled histones and analyzed the complex formation using size-exclusion chromatography (SEC). The H2a-H2b dimer and (H3-H4)_2_ tetramer used in our biochemical and biophysical studies were individually purified and assembled as reported (Jos *et al*, 2021; Tanaka *et al*, 2004; Dyer *et al*, 2004). During reconstitution, the complex mixture remained soluble at physiological salt concentration (150 mM NaCl), showing no precipitation due to non-specific interactions. As αSyn belongs to the IDP family, it eluted as a higher molecular weight protein compared to the H2a-H2b dimer in the SEC. Intriguingly, αSyn formed a ternary complex with both the H2a-H2b dimer and (H3-H4)_2_ tetramer, resulting in peak shift compared to individual components (**Figure 1B and Supplementary Figure 1**).

To further confirm αSyn association with histone assemblies, we carried out αSyn co- localization studies with H2b and H3 in SH-SY5Y cells. Only 3-7% of control SH-SY5Y cells showed αSyn in the nucleus. Previous studies have indicated an increased nuclear localization of αSyn in paraquat-treated mice, an herbicide linked with PD (Goers *et al*, 2003). So, to elevate αSyn’s nuclear level, we treated the SH-SY5Y cells with paraquat at 10 and 25 μM concentrations. Interestingly, in both control and paraquat-treated cells, αSyn co-localizes with histone H2b and H3 (**Supplementary Figure 2**). In the cellular system, individual core histones (H2a, H2b, H3, and H4) are assembled into heterodimers, H2a with H2b and H3 with H4, immediately after protein synthesis. These assembled histones are not free; they are bound by the histone chaperones and other chromatin factors that help to prevent toxic effects caused by unregulated DNA binding, leading to aggregation and interference with nuclear processes (Warren & Shechter, 2017; Gurard-Levin *et al*, 2014). Consequently, the observed αSyn co-localization with histone H2b and H3 suggests that it is possibly associated with the assembled H2a-H2b dimer and (H3-H4)_2_ tetramer in the cellular system, hinting at a role in chromatin regulation.

### αSyn has distinct binding sites for H2a-H2b dimer and (H3-H4)_2_ tetramer

To identify the αSyn region important for interactions with H2a-H2b and (H3-H4)_2_, we measured αSyn(FL) and truncated αSyn proteins binding affinity (Kd) using MicroScale Thermophoresis (MST) and Isothermal titration Calorimetry (ITC) at physiological salt concentration. MST requires Cys/Lys- labelling of target proteins for kinetic studies. Both αSyn and core histones have many Lys residues and no Cys residues except histone H3. Previously we have observed interference in binding kinetic between Lys-labeled-αSyn with the individual core histones (Jos *et al*, 2021). Therefore, we introduced a Cys-residue in the H2a C-terminal tail (G129C) and used this mutant protein to assemble H2a_G129C_- H2b dimer, which was Cys-labeled for binding studies. Upon addition of αSyn to fluorescently labeled H2a_G129C_-H2b dimer/(H3-H4)_2_ tetramer, we observed apparent changes in thermophoresis. Intriguingly, αSyn(FL) showed a robust binding affinity with H2a_G129C_-H2b dimer (Kd = 515.4 nM). Whereas αSyn(1- 131) construct showed a binding affinity of Kd = 32.4 μM, which is 64-fold lower than αSyn(FL) (**Figure 1C; i and ii**). To further validate these results, we explored the αSyn(FL) and αSyn(1-121) interaction with H2a-H2b dimers using ITC. Consistent with our MST result, αSyn(FL) binds to H2a-H2b with an affinity of Kd = 530 nM. Conversely, the αSyn(1-121) construct exhibits no binding, indicating that the αSyn(122-140) region is critical for H2a-H2b dimer interaction (**Figure 1C; iii and iv**).

Next, we conducted binding studies on αSyn’s interaction with Cys-labeled (H3-H4)_2_ tetramer. αSyn(FL) and truncated αSyn(1-103) construct lacking a complete acidic C-terminal tail showed binding affinities of Kd = 50 nM and 107 nM to (H3-H4)_2_ tetramer, respectively (**Figure 1D; i and ii**). This study suggests that, unlike the H2a-H2b dimer, the acidic stretch of αSyn is not critical for (H3-H4)_2_ tetramer binding. Similar observations were seen in the case of other histone chaperones such as NAP1, nucleoplasmin, CIA/ASF1, NO38, and SET/TAF-I*β*/INHAT, where the removal of an acidic stretch did not impair their histone chaperone or core histone-binding activities (Muto *et al*, 2007; Dutta *et al*, 2001; Daganzo *et al*, 2003; Namboodiri *et al*, 2004; Park *et al*, 2005; Umehara *et al*, 2002). The N-terminal αSyn(1-103) region upon binding to the phospholipid membrane, adopts an amphipathic α-helical secondary structure (Chandra *et al*, 2003). The electrostatic potential of this αSyn helical region shows negatively charged residues aligned on one side, suggesting a potential interaction with the (H3-H4)_2_ tetramer (**Supplementary Figure 3**). Our earlier study demonstrated that αSyn(FL) has binding affinities of Kd = 4 μM to histone H3 and H4, linker histone H1.1 with a Kd of 21 μM, H2a with a Kd of 278 μM, and for H2b with a Kd of 122 μM (Jos et al., 2021). In the current study, the αSyn showed 10- to 100-fold higher binding affinity for assembled H2a-H2b/(H3-H4)_2_ complexes than individual core and linker histones, suggesting αSyn preferential binding to histones assemblies over individual counterparts. Furthermore, truncation studies revealed two distinct binding sites in αSyn: the αSyn(1- 103) region binds to (H3-H4)_2_, while the acidic C-terminal region (121-140) is critical for the interaction with H2a-H2b.

Most canonical and variant dimer-specific H2a-H2b/H2a.Z-H2b histone chaperones share the conserved DEF/Y motif and a variable proline residue located one residue away from the motif (Wang *et al*, 2019; Kemble *et al*, 2015; Huang *et al*, 2020b). Sequence analysis of αSyn(121-140) region revealed that it shares similarities with the Swc5 DEF/Y motif, a subunit of ATP-dependent SWR chromatin remodeler that binds preferentially to canonical H2a-H2b dimer (Huang *et al*, 2020b). Unlike Swc5, which has two consecutive DEF/Y motifs, αSyn has a single DEF/Y motif and a proline residue. Further analysis reveals that the acidic αSyn(121-140) C-terminal end is well conserved across a broad range of organisms, including primates, rodents, bats, and some aquatic mammals, indicating a similar function across these species (**Figure 1E**). Together, sequence analysis has shown that the αSyn dimer-binding region is conserved across various organisms and contains a DEF/YxP motif, common to evolutionarily unrelated dimer-binding chromatin regulators.

### αSyn(121-140) binds specifically to globular domain of H2a-H2b dimer possibly via a conserved DEF/YxP motif

To delineate the structural elements, we employed a crosslinking approach to study the interaction between αSyn and H2a-H2b dimer. Cross-linking assay is a valuable technique for studying protein-protein interactions, as they covalently link two amino acid residues in protein complexes that are in proximity. This technique is widely applied in histone chaperoning studies (Bellelli *et al*, 2018; Corbeski *et al*, 2018; Chen *et al*, 2015). Here, we standardized experiments using Disuccinimidyl suberate (DSS) and 1-ethyl-3-(3-dimethylaminopropyl) carbodiimide hydrochloride (EDC) crosslinkers. DSS reacts with the primary amine group at the N-terminus of polypeptide and in the side-chain of lysine residue, whereas EDC is a carboxyl- and amine-reactive zero-length crosslinker.

The assembled H2a-H2b heterodimer consists of N-terminal flexible tails and a C-terminal globular domain (McGinty & Tan, 2015). Initial experiments were performed to investigate whether the flexible tail or the globular domain is important for αSyn interaction. Using DSS we crosslinked αSyn(FL) with H2a-H2b and tailless (H2a-H2b)TL dimers to trap their respective complexes. αSyn(FL) remains a monomer in the presence and absence of DSS. The H2a-H2b and (H2a-H2b)TL dimers run as individual bands (∼16 kDa) in the absence of DSS, but in its presence, two bands (∼16 kDa and 30 kDa) corresponding to monomer and heterodimer are seen. Subsequent titration of αSyn(FL) with H2a- H2b and (H2a-H2b)TL dimers in the presence of DSS resulted in three bands (∼16 kDa, 30 kDa, 43 kDa) corresponding to monomer, heterodimer, and their respective complexes. Interestingly, no precipitation or additional bands corresponding to higher-order complexes or aggregates were noticed even with a 2-fold excess αSyn, indicating that the αSyn interaction with the H2a-H2b dimer is specific and not driven by non-specific electrostatic interactions. Furthermore, the H2a-H2b dimer with and without N-terminal flexible tail exhibited complex formation with αSyn(FL), highlighting the importance of the globular domain over the N-terminal tail for αSyn interaction **(Figure 2A; i).**

**Figure 2:**
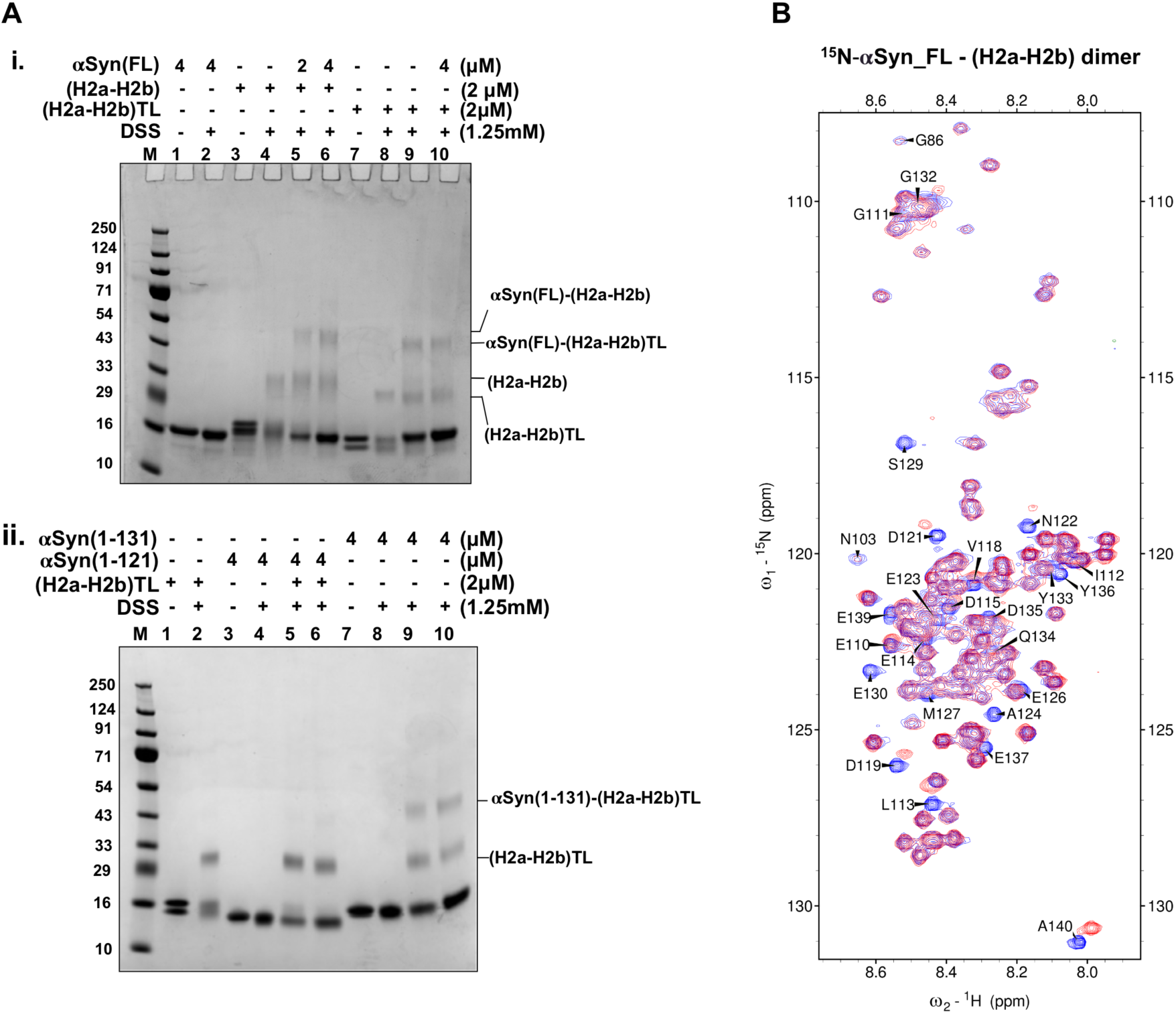
Crosslinking assay and HSQC NMR spectra. **A.** SDS-PAGE gel of DSS mediated cross- linking experiment (Lane M- protein marker along with the indicated protein concentrations at the top). i. αSyn(FL) with both (H2a-H2b) and (H2a-H2b)TL, ii. αSyn(1-131) and αSyn(1-121) with (H2a-H2b)TL. The cross-linking assays were performed in the presence of 150 mM NaCl. **B.** HSQC NMR spectra of ^15^N-labeled αSyn(FL) in the absence (blue) and presence (red) of (H2a-H2b) dimer (1:1 ratio) showed significant chemical shift perturbations for αSyn C-terminal (120-140) residues.

Subsequently, we characterized the αSyn region that specifically associates with (H2a-H2b)TL dimers. The αSyn(1-131) showed binding, whereas the αSyn(1-121) did not bind to (H2a-H2b)TL dimers (**Figure 2A; ii**). This study further reiterated our MST and ITC data and unambiguously demonstrated that the αSyn acidic C-terminal tail (121-140) is essential for H2a-H2b interaction, and its removal abolishes complex formation. To validate the above findings, we custom synthesized αSyn(121-140) peptide and analyzed its interaction with (H2a-H2b)TL using DSS and EDC crosslinkers. The αSyn(121-140) region that lacks lysine residue and has an NH2-group only at the polypeptide N-terminal, showed a minor shift in the dimer band with DSS and a clear shift with EDC (**Supplementary Figure 4**).

To corroborate our findings further, we performed NMR chemical shift perturbation mapping and compared the ^1^H-^15^N heteronuclear single-quantum coherence (HSQC) spectra of the uniformly ^15^N-labeled αSyn(FL) in the absence and presence of unlabeled H2a-H2b dimer. The results indicated that the N-terminal part of αSyn(1-120) remains largely unaffected, whereas a change in peak intensity and chemical shift was observed at the αSyn C-terminal end. Specifically, the residues in the αSyn(121- 140) region underwent structural reorganization upon binding to H2a-H2b, which correlates with the cross-linking and biophysical data (**Figure 2B**).

### αSyn(121-140) with single-chain H2a-H2b dimer complex structure

To elucidate the mechanism of H2a-H2b dimer recognition by αSyn, we determined the crystal structure of αSyn(121-140) in complex with the H2a-H2b dimer at 1.72 Å resolution (**Figure 3A and Table 1**). The rationale for using ScH2a-H2b for structural studies has been well established (Wang *et al*, 2019; Zhou *et al*, 2008; Mao *et al*, 2014; Liang *et al*, 2016). The αSyn(121-140)- ScH2a-H2b complex structure was solved by molecular replacement using PDB ID: 6W4L as a search model, and the refined final model has Rwork/Rfree of 19.0/21.9. There were two molecules in the asymmetric unit, which is superimposed with an r.m.s.d value of 0.43 Å. The αSyn(136-140) region from both molecules in the asymmetric unit overlaps but runs in an opposing direction (**Figure 3B**). Intriguingly, αSyn(121-140) adopts two binding modes (BM-1 and BM-2) when interacting with the dimer. Biophysical studies on H2a_G129C_-H2b dimer with αSyn(121-140) peptide using MST supported our structural data, revealing two binding sites with Kd values of 0.5 and 2.6 *μ*M (**Supplementary Figure 5A**). For BM-1 and BM-2, the electron density is missing for residues 121-130 and 121-127, respectively. Overall, the electron density for BM1 is relatively weak and localized, whereas BM2 shows good density for both main- and side-chains. However, in both binding modes, the electron density gradually wanes after the Y133 residue as we proceed toward the N-terminal region (**Supplementary Figure 5B and 5C**).

**Figure 3.**
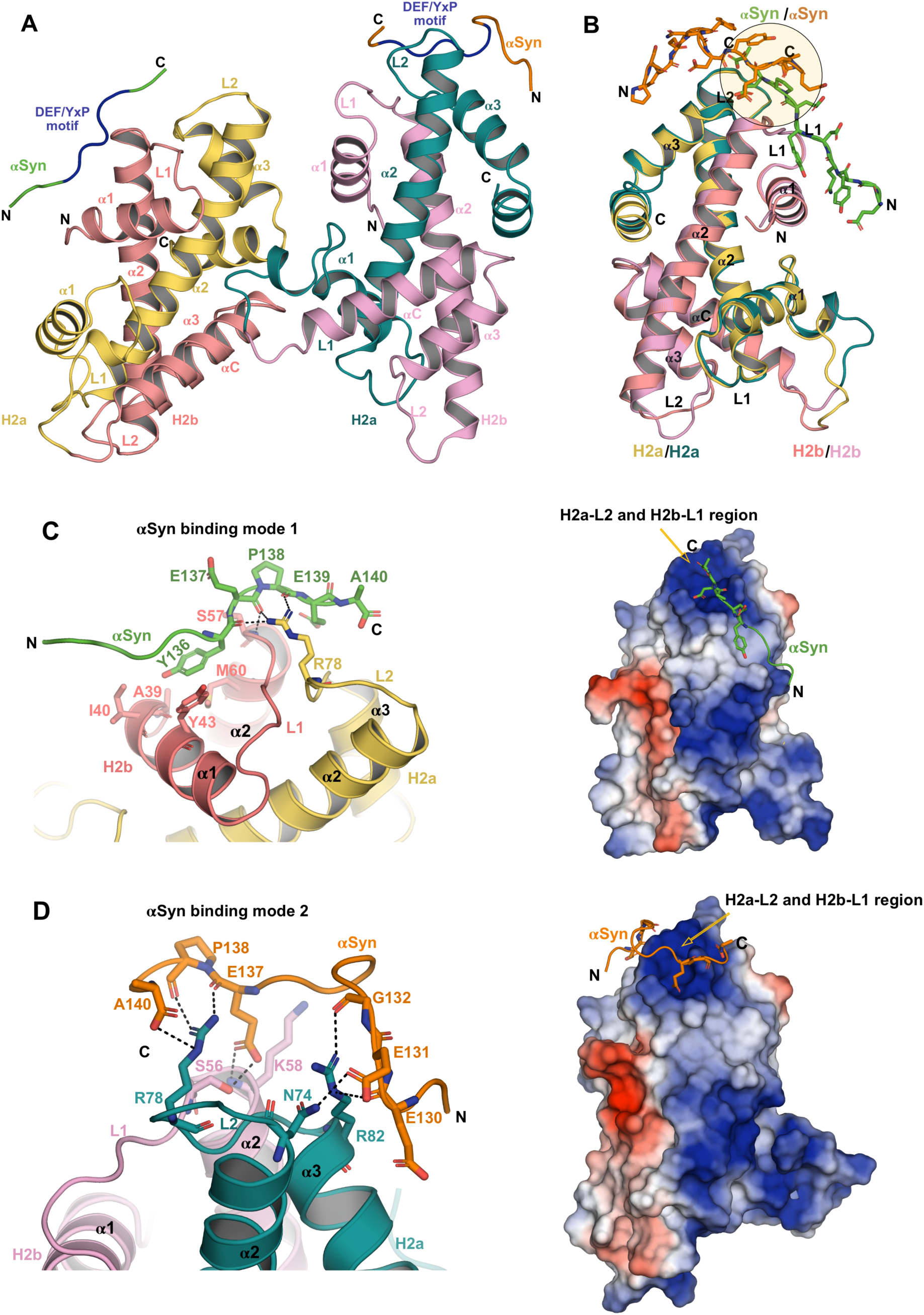
Structural basis of αSyn with H2a-H2b complex. **A.** The overall structure of αSyn(121- 140)-ScH2a-H2b complex is shown in cartoon representation. The asymmetric unit contains two molecules, and αSyn has different binding modes for each unit of ScH2a-H2b. The DEF/YxP motif is indicated in blue. **B.** Superimposition of αSyn(121-140)-ScH2a-H2b complex within asymmetric unit shows that αSyn(136-140) overlaps and runs in opposing directions and is highlighted in a circle. **C-D.** αSyn binding mode-1 and 2; close-up view of residues involved in the αSyn-scH2a-H2b interface interactions are shown in stick representation (left). The interface between αSyn and ScH2a-H2b for both binding modes is shown in the electrostatic surface representation (right).

**Table 1:** Data collection and refinement statistics.

| | αSyn(121-140)–ScH2a-H2b dimer<br>complex structure (PDB code: 8ZVY) |
| --- | --- |
| Data collection and processing |  |
| Space group | P2<sub>1</sub>2<sub>1</sub>2<sub>1</sub> |
| Cell dimension |  |
| a, b, c (Å) | a=61.0, b=68.2, c=102.5 |
| $\alpha$, $\beta$, $\gamma$ (°) | $\alpha=\beta=\gamma=90^\circ$ |
| R-merge (%) | 3.0 (85.7) |
| Resolution range (Å) | 68.24- 1.72 (1.75 - 1.72) |
| No. of unique reflections | 46279 (2444) |
| Completeness (%) | 100 (99.9) |
| CC<sub>1/2</sub> | 100 (87.9) |
| Multiplicity | 12.1 (12.1) |
| I/$\sigma$(I) | 37.1 (3.2) |
| Structure refinement |  |
| R<sub>work</sub> | 0.1907 |
| R<sub>free</sub> | 0.2205 |
| No. of non-H atoms | 3284 |
| ScH2bH2a chains/atoms | 3054 |
| Water | 228 |
| ligand | 2 |
| R.m.s deviations |  |
| Bonds lengths (Å) | 0.006 |
| Angles (°) | 0.89 |
| Ramachandran |  |
| Favored (%) | 98.41 |
| Allowed (%) | 1.59 |
| Outlier (%) | 0 |
| Average B-factor | 47.6 |
| r.m.s., root mean square.<br>Highest resolution shell is shown in parenthesis. One crystal was used for the data.<br><sup>c</sup>R<sub>free</sub> is the R factor for a subset of 5% of the reflections that were omitted from refinement. |  |

### αSyn DEF/YxP motif interacts extensively with the L2-L1 region of H2a-H2b dimer

Structural analyses showed that the αSyn DEF/YxP motif in both binding modes interacts with the H2a-H2b dimer in the H2a-L2 and H2b-L1 loop regions and makes extensive contact with the H2a- R78 residue. In BM1, the buried surface area between αSyn and H2a-H2b dimer is 450.4 Å. In the case of BM1, intermolecular interaction between αSyn with H2a-H2b is localized, involving the main-chain CO of E137 and P138, as well as both the main- and side-chain of Y136; thus, only these residues showed reasonable density. The αSyn Y136 and E137 form a hydrogen bond with H2a-R78 NH2 and P138 CO with H2a-R78 NH1, respectively, while the main chain of αSyn E137 CO forms a hydrogen bond with H2b-S57 NH. Additionally, the side chain of αSyn Y136 is buried in the shallow hydrophobic pocket surrounded by A39, I40, and Y43 in the H2b-α1 helix and M60 in the H2b-α2 helix, and forms a **π**-**π** interaction with H2b-Y43 (**Figure 3C**). In BM-2, the buried surface area between αSyn and H2a- H2b dimer is 382.8 Å^2^. The main-chain CO of αSyn E137 and P138 form hydrogen bonds with H2a R78 side-chain NH1 and NH2, while the αSyn COOH group at the C-terminal end forms salt bridges with H2a R78 NE. Similarly, the main-chain CO of αSyn G132 and E130 form hydrogen bonds with the side-chain of H2a R82 NE and NH1, respectively. Additionally, the side chain of αSyn E131 OE1 forms a hydrogen bond with H2a N74 ND2. Furthermore, the side-chain of αSyn E137 OE1 and OE2 form hydrogen bonds with main-chain K58 N and with side-chain H2b S56 OG in the H2b-L1 loop (**Figure 3D**). Overall, the αSyn DEF/Y motif, together with a conserved P138 residue, interacts extensively with the L2-L1 region of the H2a-H2b dimer, and their interactions are stabilized through an electrostatic anchor flanked by polar and hydrophobic interactions.

### Dimer recognition by αSyn overlaps with other histone chaperones

Overlay of αSyn(121-140)–ScH2a-H2b dimer with canonical H2a-H2b specific Spt16 (Kemble *et al*, 2015) and Swc5 (Huang *et al*, 2020b), and H2a.Z-H2b variant specific YL1 (Latrick *et al*, 2016; Liang *et al*, 2016), Chz1 (Zhou *et al*, 2008), and Anp32e (Mao *et al*, 2014; Obri *et al*, 2014) chaperone complex structures reveals that they all have overlapping dimer recognition site. In αSyn-BM1, the positions of E137 and Y136 residues are conserved across other dimer-specific histone chaperone structures (**Figure 4A and 3C**). Mutational studies of Swc5-F29 and Spt16-Y972 (Huang *et al*, 2020b; Kemble *et al*, 2015), corresponding to αSyn-Y136 residue, showed a substantial reduction in binding affinity due to the loss of specific hydrophobic contact. In the case of BM2, although the position of αSyn E137 is conserved, the αSyn interaction with dimer is rather distinct, with its COOH group at the C-terminal end also involved in capping H2a-R78 residue (**Figure 4B and 3D**). This study suggests that although negative charges of αSyn are crucial for histone binding, specific recognition via an aromatic anchor together with the conserved P138 residue is important for interaction with H2a-H2b dimer.

**Figure 4:**
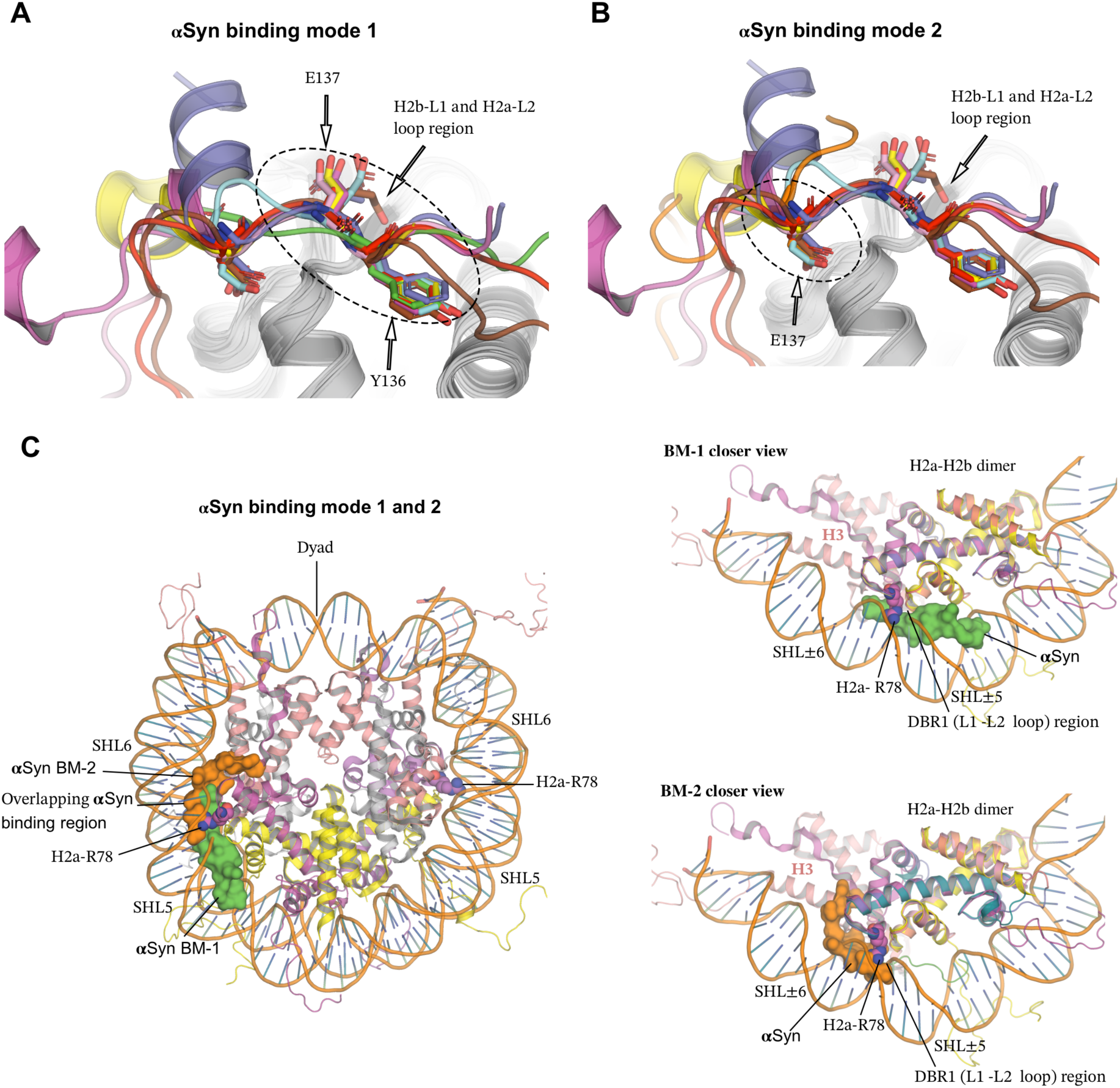
αSyn binding region of H2a-H2b overlaps with other dimer-specific histone chaperones. **A-B.** Superimposition of αSyn BM1 and BM2 with other known dimer-specific chaperone structures. αSyn-BM1 (green), αSyn-BM2 (orange), Anp32e (PDB: 4CAY, pink), YL1 (PDB: 5CHL, red), YL1 (PDB: 5CHL, red), Swc5 (PDB: 6KBB, purple), Chz1 (PDB: 6AE8, yellow), Spt16 (PDB: 4WNN, cyan), YL1 (PDB: 5FUG, brown), Spt16 (PDB: 8I17, blue) and H2a-H2b/H2a.Z-H2b dimer (gray). In BM1, the position of Y136 and E135 is conserved with other histone chaperones, whereas in BM2, the position of E137 is conserved, indicated in a dotted circle. **C.** Superimposition of αSyn(121-140)-ScH2a- H2b complex structure with the NCP structure (PDB ID: 1KX5). The H2a-L2 loop R78 residue anchored to a minor groove of the DNA at SHL±5.5 in the nucleosome shown in spheres. The αSyn(136-140) region in both binding modes (BM-1 and BM-2) overlaps, exclusively binds to DBR1 (H2a-L2 and H2b- L1 loop), and caps conserved H2a-R78 residue (left). In BM1, αSyn peptide clashed with the DNA-binding site of H2a-H2b in the nucleosome (right-top). In BM2, αSyn peptide clashed with both DNA-binding sites of H2a-H2b and competes for the H3 binding site (right-bottom).

The H2a-H2b dimer in the assembled nucleosome protects its entry/exit site by interacting with nucleosomal DNA on one side and (H3-H4)_2_ tetramer on the other. There are three main nucleosomal DNA interaction interfaces on H2a-H2b referred to as ‘DNA binding region (DBR)’: the L1-L2 binding sites at superhelix locations (SHL) ±5.5 (DBR-1) and ±3.5 (DBR-3) flanking the α1-α1 middle binding site at SHL ±4.5 (DBR-2) (Huang *et al*, 2020a). Superimposition of αSyn(121-140)–ScH2a-H2b dimer with nucleosome core particle (NCP; PDB: 1KX5) shows that αSyn has two distinct interaction sites within the nucleosome. In BM1, αSyn exploits the nucleosomal DNA binding surface, while in BM-2, αSyn interacts with both the DNA-binding surface and competes for the H3 interaction surface with the H2a-H2b dimer (**Figure 4C**). Notably, in both binding modes (BM-1 and BM-2), the αSyn(136-140) region overlaps and exclusively targets DBR1, capping the side chain of the conserved H2a-R78 residue, which otherwise anchors to the minor groove of DNA at SHL±5.5 within the nucleosome.

Electromobility shift assay (EMSA) shows no change in the NCP mobility with increasing concentration of αSyn(FL), suggesting no binding between the two molecules (**Supplementary Figure 6**). This study indicates that αSyn binding sites on histone H2a-H2b/(H3-H4)_2_ are inaccessible when assembled within the NCP. Additionally, our earlier study indicated that αSyn’s interaction with dsDNA is weak and non-specific. Collectively, the overlapping dimer recognition site of αSyn and other histone chaperones denotes that αSyn might act as a gatekeeper during the histone eviction/deposition step in the nucleosome assembly/disassembly process.

## DISCUSSION

The nuclear αSyn role is coupled with gene expression (Kontopoulos *et al*, 2006), DNA repair (Schaser *et al*, 2019), and transcriptional regulation (Pinho *et al*, 2019; Paiva *et al*, 2017; Davidi *et al*, 2020). Under pathological conditions, αSyn exhibits excessive nuclear localization, which adversely impacts gene expression in vulnerable neurons through transcriptional dysregulation, altered splicing, and compromised DNA repair processes (Paiva *et al*, 2017; Sepe *et al*, 2016; Grünblatt *et al*, 2004; Kontopoulos *et al*, 2006). Nonetheless, the molecular basis of how αSyn regulates chromatin functions under physiological conditions—and how these are altered in pathological states—remains poorly understood. Even the precise nuclear function of αSyn remains unclear. In this study, we discovered the crucial nuclear physiological roles of αSyn as a histone chaperone and showed its potential role in nucleosome assembly and disassembly. Recently, the involvement of histone chaperone Anp32e in memory formation, transcription, and dendritic morphology in neurons was reported (Stefanelli *et al*, 2021), highlighting the functional significance of our findings.

The brain development in vertebrates necessitates a complex interplay between developmentally dynamic alternative splicing and gene expression (Mazin *et al*, 2021). Both transcription and alternative splicing (AS) are coupled processes (de la Mata *et al*, 2003), and studies indicate that neuronal cells expand their transcription diversity by AS of precursor mRNA (pre-mRNA) (Paiva *et al*, 2017). AS is highly conserved and prevalent, contributing significantly to the functional complexity of the nervous system. Studies suggest that the balance of AS can be modulated by the availability of histones during transcriptional elongation by RNA polymerase II (Pol II). A fast transcription elongation rate favors exon skipping, whereas a slow rate allows recognition of weak splice sites. Therefore, establishing an optimal elongation rate is a prerequisite for normal co-transcriptional pre-mRNA splicing (Dujardin *et al*, 2014; Fong *et al*, 2014). Even modest changes in the elongation rate, either increase or decrease, can have substantial effects on splicing, a phenomenon widely documented in cancer and other diseases. Intriguingly, issues with transcription dysregulation and defects in AS are also reported across various neuropsychiatric and neurodegenerative diseases, including PD (Fu *et al*, 2013; Li *et al*, 2021; La Cognata *et al*, 2015; Licatalosi & Darnell, 2006; Nikom & Zheng, 2023; Tollervey *et al*, 2011). Nonetheless, how αSyn-induced transcriptional and splicing deregulation occurs and the underlying nuclear pathological mechanism remains unclear.

Drawing on our research and that of others, we propose a model outlining nuclear-localized αSyn’s role in physio-pathological conditions (**Figure 5**). During eukaryotes gene transcription, the nucleosome must disassemble ahead and reassemble behind Pol II as elongation progresses. This highly regulated process necessitates physical interaction between histone chaperones and chromatin assembly factors to ensure a timely and accurate supply of histones during the nucleosome assembly and disassembly (Groth *et al*, 2007; Venkatesh & Workman, 2015). Noticeably, decreasing canonical histone availability can accelerate the Pol II elongation rate, leading to splicing defects (Jimeno-González *et al*, 2015; Prado *et al*, 2017; Dujardin *et al*, 2014; Murillo-Pineda *et al*, 2014; Hu *et al*, 2014; Fong *et al*, 2014). In this study, we demonstrated that αSyn binds to the assembled histone H2a-H2b dimer and (H3-H4)_2_ tetramer with high affinity and specificity. Furthermore, our structural study also shows that αSyn and other dimer-binding chromatin regulators share a common overlapping histone recognition site, highlighting its potential role in chromatin dynamics. Based on these findings, we contemplate that excessive nuclear accumulation of αSyn under pathological conditions might deplete the available histone pool during transcription. This depletion could lead to aberrant splicing, shifting susceptible neuronal cells from producing physiologically relevant isoforms to generating inactive or aberrant protein isoforms, thereby depriving neurons of vital transcripts.

**Figure 5:**
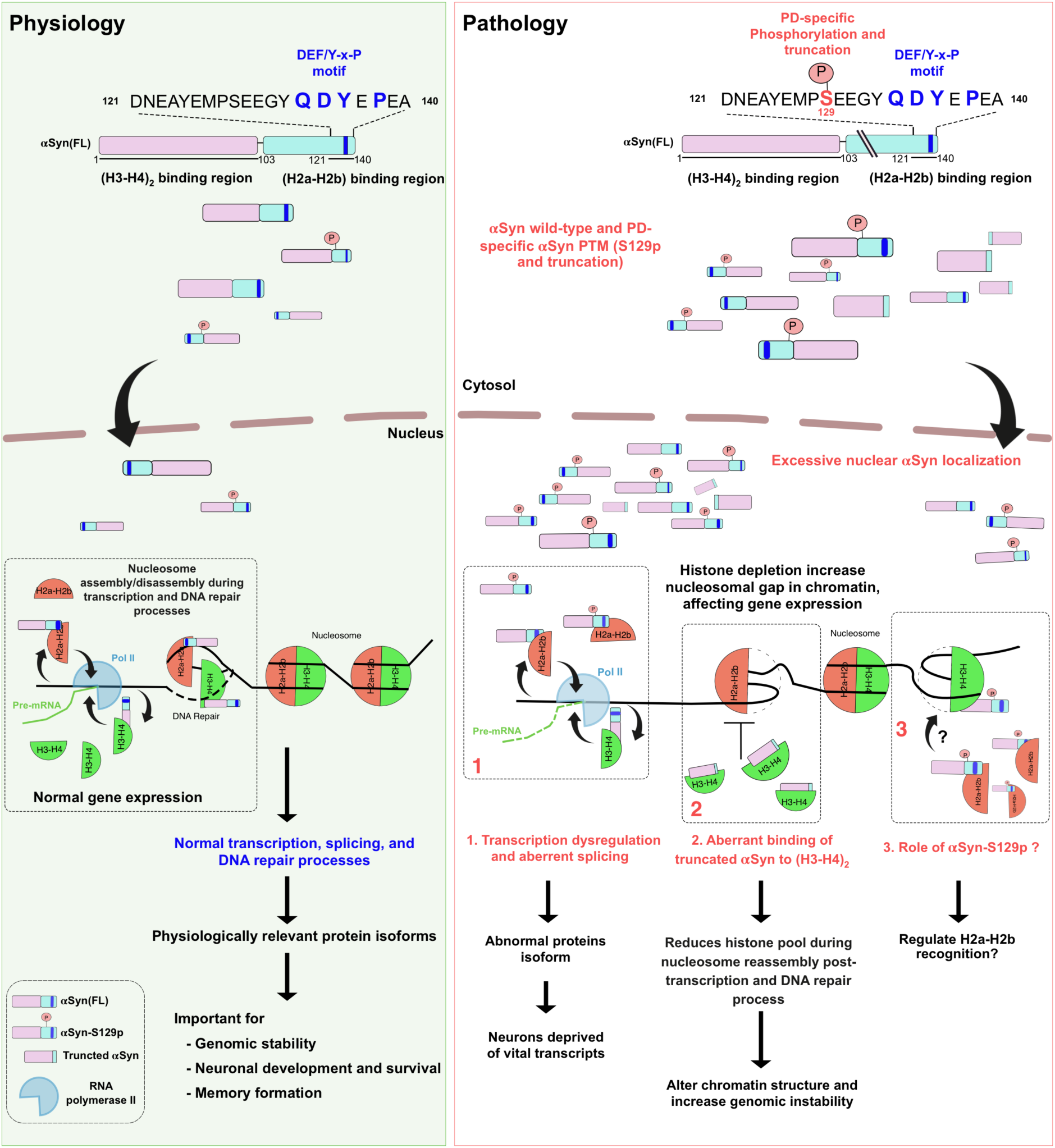
Model of nuclear αSyn role in physio-pathological conditions. Under physiological conditions, the nuclear-localized αSyn possibly regulates nucleosome assembly/disassembly during transcription and DNA repair, which is essential for normal gene expression. Conversely, excessive nuclear αSyn localization depletes the histones pool, increasing the nucleosomal gap and adversely affecting gene expression under pathological conditions.

Second, studies have shown that DNA damage foci and DNA breaks significantly increase during aging (Hu *et al*, 2014; Madabhushi *et al*, 2014). This DNA damage could progressively alter chromatin structure, impacting genome integrity and stability, thereby affecting gene expression patterns as aging progresses. Cells may use transcription machinery to monitor DNA integrity and activate DNA damage signaling (Ljungman & Lane, 2004). Since blockage in transcription due to DNA lesions can trigger apoptosis, it is crucial for cells to quickly resolve these blockages and restore RNA synthesis. Studies show that the PD brain tissue from the *substantia nigra* has altered αSyn splice variant expression; it exhibits higher levels of C-terminally truncated αSyn transcripts (SNCA-112 and SNCA-98) compared to normal conditions (Cardo *et al*, 2014; McLean *et al*, 2012). Our studies strongly suggest that these pathological αSyn splice variants do not interact with the H2a-H2b dimer; instead, they might aberrantly bind (H3-H4)_2_ tetramer, which could alter the histone availability during the nucleosome reassembly post-transcription and DNA repair. Specifically, subtle changes in chromatin caused by a deficit in the pool of available histones have deleterious consequences on genome integrity.

Third, the acidic αSyn C-terminus serves as a central hub for protein-protein interactions (Parra- Rivas *et al*, 2023) and harbors various post-translation modifications (PTM) sites (Manzanza *et al*, 2021). Among the αSyn PTMs, S129-phosphorylation holds physio-pathological significance; only 4% of αSyn is phosphorylated at the S129 position in the normal brain (Ramalingam *et al*, 2023; Anderson *et al*, 2006), compared to 90% under pathological conditions (Fujiwara *et al*, 2002). The αSyn(S129A) mutant, which blocks phosphorylation, forms cytoplasmic inclusion, suggesting that this PTM acts as a molecular switch controlling αSyn nuclear localization (Gonçalves & Outeiro, 2013). Likewise, toxic familial PD mutants (G51D, E46K, A30P, and A53T) exhibiting varying aggregation propensity (Mehra *et al*, 2019), share a common characteristic of enhanced nuclear accumulation (Fares *et al*, 2014; Mbefo *et al*, 2015; Gonçalves & Outeiro, 2013). In various biological contexts, adding or removing a dianionic phosphate group often alters the protein’s structural properties and modulates protein-protein interactions (Nishi *et al*, 2014; Bah *et al*, 2015). Our study has revealed that the DEF/YxP motif at the αSyn C-terminal end is critical for anchoring the H2a-H2b dimer. Given that the S129-phosphorylation site is adjacent to the DEF/YxP motif, this PTM might induce conformational changes at the C-terminus, thus potentially regulating its interaction with the H2a-H2b dimer. While studies linking αSyn-S129 phosphorylation with LB formation have been extensively explored, our finding suggests that this PTM might also significantly impact its interaction with histone assemblies. Therefore, investigating how PD-specific αSyn-S129 phosphorylation regulates H2a-H2b dimer binding could provide insights into its nuclear physio-pathological role. In conclusion, future studies aimed at molecular-level understanding of αSyn’s role in chromatin regulation—including gene expression, transcription, and DNA repair—are crucial for gaining detailed insights into its physio-pathological roles.

## Materials and Methods

### Cloning, expression, and purification of human core histone proteins

Human full-length core histones H2a, H2b, H3, and H4 constructs, along with the N-terminal tail truncated core histones H2a and H2b, were cloned, expressed, and purified as previously described (Tanaka *et al*, 2004; Jos *et al*, 2021). In brief, the full-length (FL) and N-terminal tail less (TL) core histones with (His)_6_-tag at N-terminus were expressed in *E.coli* expression strains BL21(DE3) (histone H2a, H2b, H3, H2aTL, and H2bTL) or JM109(DE3) (histone H4) cells as inclusion body and purified under a denaturing condition. The purified histones were lyophilized and stored at -80℃ until further use. H2a_G129C_ mutation was generated for labeling during MST studies and purified using standard histone H2a purification steps.

### Cloning, expression, and purification of αSyn proteins

αSyn(FL) was used as a template to create C-terminal truncated αSyn(1-131), αSyn(1-121), and αSyn(1-103) constructs and sub-cloned into pET28a vector (Novagen) at NdeI and BamHI restriction sites. The human αSyn(FL) was expressed and purified from periplasmic space as previously described (Jos *et al*, 2021). Whereas αSyn truncation constructs with (His)_6_-tag at N-terminus were transformed in *E.coli* BL21(DE3) cells, induced with 0.5 mM isopropyl β-d-1-thiogalactopyranoside (IPTG) with post-induction at 37°C for 5 hours. The cells were lysed and centrifuged, and the supernatant was loaded onto the affinity chromatography using IMAC FF 5ml column (GE Healthcare), followed by ion-exchange purification using the Q FF column (GE Healthcare). The His-tag was cleaved with thrombin digestion overnight, and the sample was then purified using a Superdex 75 HiLoad 16/600 gel filtration column (GE Healthcare). All purified αSyn constructs were lyophilized and stored at -80℃ until further use.

### Reconstitution of the H2a-H2b, (H3-H4)_2_ and αSyn with histone assembly complexes

Reconstituted H2a-H2b and (H2a-H2b)TL dimers, and (H3-H4)_2_ tetramer using previously published methods (Dyer *et al*, 2004). The purified individual core histones were dissolved in unfolding buffer (6 M Guanidium chloride, 20 mM Tris-HCl (pH 7.5), and 5 mM DTT), mixed at 1:1 stoichiometry, and dialyzed overnight against the refolding buffer containing 20 mM Tris-HCl pH 8.0, 150 mM NaCl, 1 mM EDTA, and 5 mM β-mercaptoethanol (β-ME). The reconstituted histone assemblies were purified by loading into size-exclusion chromatography (SEC) using Hiload 16/60 Superdex 200 (GE Healthcare). Then, αSyn(FL)-(H2a-H2b) and αSyn(FL)-(H3-H4)_2_ complexes were assembled by mixing αSyn(FL) with H2a-H2b in 1:1.1 stoichiometry, likewise αSyn(FL) with (H3-H4)_2_ tetramer in 1.1:1 stoichiometry and analyzed using SEC for ternary complex formation. The individual components, αSyn(FL), H2a-H2b dimer, (H3-H4)_2_ tetramer, and respective αSyn-histone assembly complexes were injected independently to SEC equilibrated in 20 mM Tris pH 8.0, 150 mM NaCl, and 2 mM β-ME buffer.

### MicroScale thermophoresis (MST)

The MST experiments were performed according to the NanoTemper technologies protocol, and affinities were calculated using the Monolith NT.115 (Red/blue) instrument (NanoTemper Technologies GmbH, Munich, Germany). For MST experiments, (H2a_G129C_-H2b) dimer and (H3-H4)_2_ tetramer were labeled using cysteine reactive Monolith NT™ Protein Labeling Kit RED-MALEIMIDE (NanoTemper Technologies) for interaction studies of various αSyn constructs. The final concentration of NT-647 labeled (H2a_G129C_-H2b) dimer and (H3-H4)_2_ tetramer were 150 nM each. The above-labeled concentrations were chosen based on fluorescence intensities from the pretest assay setup using MO.Control 1.5.3 software (NanoTemper Technologies GmbH). For this study, all the samples were prepared as previously described (Jos *et al*, 2021), and data was acquired at 25°C using 90% LED power and 40 % MST power. The MST setup for each binding affinity was analyzed separately and the *k_d_* values were determined with MO.Affinity Analysis 2.2.7 software (NanoTemper Technologies GmbH).

### Isothermal Titration Calorimetry (ITC)

The purified αSyn(FL), αSyn(121), and H2a-H2b dimer proteins were buffer exchanged with PBS buffer. Then, ITC experiments were performed with the above sample using a MicroCal ITC 200 instrument at 25°C on high feedback mode with a stirring speed of 800 rpm and a filter period of 5 s. 200 μL of 50 μM H2a-H2b dimer was titrated with 350 μM of αSyn(FL) and αSyn(121). The titration experiments were performed in 16 injections with 2.5 μL per injection and 150 second intervals between each injection. A control experiment was also performed by replacing H2a-H2b dimer with buffer to account for the heat of dilution and subtracted from the titration data. The resulting isotherms were fitted using one site model by varying the parameters N, Ka, and ΔH.

### Crosslinking assay

The purified αSyn(FL), αSyn(1-131), αSyn(1-121), and assembled and purified H2a-H2b and (H2a-H2b)TL dimers were used for crosslinking studies. Additionally, αSyn(121-140) peptide synthesized to >95% purity from GL Biochem (Shanghai, China) was used in this assay. The crosslinkers, DSS and EDC (G-Biosciences), were dissolved in DMSO at stock concentrations of 12.5 mM and 120 mM, respectively. All proteins used for cross-linking studies were buffer exchanged with 20 mM HEPES pH 6.5 buffer before the experiments. Initially, for all assay combinations, histone complexes at a concentration of 2 μM and 2 – 4 μM of αSyn proteins were mixed and incubated for 15 minutes at room temperature. Subsequently, corresponding crosslinkers were added to the protein mixtures at a final concentration of 1.25 mM for DSS and 12 mM for EDC. The reaction mixture was further incubated at room temperature for 20 minutes. The reaction was quenched using 1.0 M Tris pH 7.5 buffer with a final concentration of 50 mM in the assay, and samples were analyzed by NuPAGE™ 4 to 12% Bis-Tris 1.0 mm Mini Protein Gels (Invitrogen) in 1X MES pH 6.5 running buffer. The protein bands were visualized by Coomassie Brilliant Blue staining.

### Nuclear magnetic resonance Spectroscopy

Uniformly ^15^N-isotopically labeled αSyn(FL) was produced and stored as reported previously (Jos *et al*, 2021). The ^1^H-^15^N HSQC spectra were collected at 290 K with 2048 points and 256 t1 increments, 8 scans per t1 point, and a 1.5 s recycle delay with sweep widths of 7211 Hz (^1^H) and 1702 Hz (^15^N). The experiments were performed with 100 μM of αSyn(FL) in PBS supplemented with 10% (v/v) D_2_O on a 600 MHz Bruker Avance III HD spectrometer equipped with a cryoprobe. The data were processed with Bruker TopSpin software and analyzed with NMRFAM-SPARKY (Lee *et al*, 2015). Backbone amide resonance assignment was performed based on the reported structure (BMRB 19337) (Kang *et al*, 2013). In the interaction study with H2a-H2b dimer, a required volume of about 250 μM of ^15^N-αSyn(FL) and unlabeled H2a-H2b dimer was mixed to obtain a final solution with 100 μM of αSyn(FL) and H2a-H2b dimer.

### Crystallization and Data Collection

The single-chain Xenopus H2a-H2b dimer (ScH2a-H2b) construct was provided by Dr. David Shechter, Department of Biochemistry, Albert Einstein College of Medicine, USA. The expression and purification of the ScH2a-H2b dimer were performed as previously described (Warren *et al*, 2020). The purified ScH2a-H2b was mixed with αSyn(121-140) peptide at 1: 2 molar ratio in the 25 mM Tris pH 8.0 buffer, 0.5 mM EDTA, 1.0 M NaCl. Then, the sample was gradually diluted using 25 mM Tris pH 8.0, 0.5 mM EDTA, and 1.0 mM NDSB-256 to achieve a final salt concentration of 375 mM NaCl. The resulting complex was concentrated to 11 mg/ml, and the crystal was obtained in a couple of days by sitting-drop vapor diffusion method at 18°C by mixing equal amounts of complex and reservoir solution containing 100 mM Tris pH 8.0 and 10% PEG 8000. The crystal was optimized for cryoprotection using an in-house X-ray diffractometer at NIMHANS, Bangalore. The final dataset was collected at the XRD2 beamline at the Elettra synchrotron-radiation source, Trieste, Italy, using a Dectris PILATUS 6M detector at 100 K by cryoprotecting the crystals in reservoir solution supplemented with 20% glycerol. The data sets were indexed and scaled using iMOSFLM and AIMLESS from the CCP4 program package (Agirre *et al*, 2023).

### Structure determination and refinement

The structure was determined by molecular replacement method using PHASER with PDB ID: 6W4L as a search model. Model building and structure refinement were performed using REFMAC5, Phenix, and COOT (Murshudov *et al*, 2011; Emsley & Cowtan, 2004; Liebschner *et al*, 2019). From the beginning of the refinement, 5% of the total reflections were set aside to monitor the Rfree values. PyMOL program was used to visualize and produce figures (DeLano W.L, 2002).

### NCP assembly and electrophoretic mobility shift assay (EMSA)

NCPs assembled with recombinant human histones octamer and 145bp 601L-DNA fragments (Chua *et al*, 2012). NCP interaction was carried out by varying αSyn(FL) concentration from 1:1 to 1:4 ratio. The sample was incubated for 20 mins in buffer containing 20 mM Tris-HCl pH 8.0, 75 mM NaCl, and 2 mM β-ME before performing an EMSA using 6% Native-PAGE and analyzed gel using ethidium bromide staining.

### Immunocytochemistry

For the nuclear co-localization study, SH-SY5Y neuronal cells (ECACC; Sigma-Aldrich) were cultured in DMEM media (Gibco) with 15% FBS (Gibco) and 1% PSN (Gibco). The cells were seeded onto 24-well plates (Corning Incorporated CoStar) with 2% gelatin (Sigma, #G1890)-coated glass coverslips at a concentration of 15,000 cells/cm². After 48 h of seeding, the cells were treated with paraquat (stock of 200 mM in DMSO) diluted with media to 10µM and 25µM stock for 24 hours. The control cells were also treated with an equal volume of DMSO. Then, cells were washed 3 times with PBS and fixed with Karnovsky’s fixative buffer for 1 hour at room temperature (Valappil *et al*, 2022). The fixed cells were washed three times with PBS and then incubated with primary antibodies, dilution of 1:150 for anti-αSyn (Cloud Clone, #PAB222Hu01) and 1:500 anti-H3 (Invitrogen, # AHO1432) or 1:1000 anti-H2b (Invitrogen, #MA5-31410), in an incubation buffer (0.1% Saponin, 0.1% tween and 5% FBS in PBS) overnight at 4°C. Then, after washing, secondary antibodies were used with a dilution of 1:1500 anti-mouse Alexa Fluor 555 (Invitrogen) and 1:1500 anti-rabbit Alexa Fluor 488 (Invitrogen) in incubation buffer for 90 mins at room temperature. The cells were then stained with DAPI, mounted on a slide, and imaged using a confocal microscope with 40x (oil) immersion objective (Zeiss LSM 980, Carl Zeiss). The images were analyzed using Fiji (NIH, USA).

### Statistics and reproducibility

ITC and MST experiments were performed in triplicates and analyzed using the respective software. One-way ANOVA was performed to assess the significant difference between control and treated cells for the subcellular localization studies, with p < 0.1 (F value = 5.43). All graphs presented in the manuscript and the statistical analyses of the confocal data were carried out using GraphPad Prism 10.1 software.

## Data availability

Coordinates and structure factors of the αSyn(121-140)–ScH2a-H2b dimer complex structure have been deposited in the Protein Data Bank under accession code 8ZVY and are publicly available as of the publication date. Microscopy data and protein constructs reported in this paper are available from the lead contact upon request.

## Supplementary data

Supplementary Figure 1 - 6.

## Acknowledgments

This work was supported by a SERB-ECR grant to PS (ECR/2018/002219) and NIMHANS intramural support. SJ, AK fellowships are supported by ICMR (ICMR-SRF (ID: 2021-8645/Genomics-BMS)) and DBT (DBT (ID: DBT/2021-22/NIMHANS/1685)) respectively. SN thanks ICMR (IIRP-2023-0084) grant for the support. The in-house X-ray diffraction facility at NIMHANS was used for initial crystal screening and confocal imaging acquisition was carried out using the NIMHANS core facility. Thanks to Ashok Sridhar and Girish P Waghmare at NIMHANS for their support in in-house X-ray data collection and confocal image acquisition. We thank the Department of Science & Technology (DST), Government of India (DST- FIST: SR/FST/LS-I/2017(C)) for the infrastructure grant. Special thanks to XRD2 beamline, Elettra synchrotron staff Dr. Raghurama P Hegde for data collection and it was possible through grant-in-aid from the DST, India, vide grant number DSTO-1668. Hemanga Gogoi supported biochemical studies. The NMR data were acquired at the National Center for Biological Science-Tata Institute of Fundamental Research NMR Facility. SP dedicates this work to his Ph.D. mentors, Prof. Tilman Schirmer and Dr. Zora-Housley Morcovick at Biozentrum, University of Basel, Switzerland.

## Author contributions

SP conceived the project, designed the experiments, solved the crystal structure, and data analysis. SJ performed cloning, purification, biophysical studies, crystallization, data collection, and confocal imaging. AK performed purification, crystallization, and data collection. TKP and NK supported NMR and ITC experiments. Shylaja Parthasarathi, SJ, and SN performed cellular studies and confocal imaging. BP for scientific inputs and structure solution. SJ, NK, and SN supported data analysis and manuscript preparation. SP wrote the original draft.

