## Supplementary Figure 1-6 for "Structural and functional insights into nuclear role of Parkinson’s Disease-associated α-Synuclein"

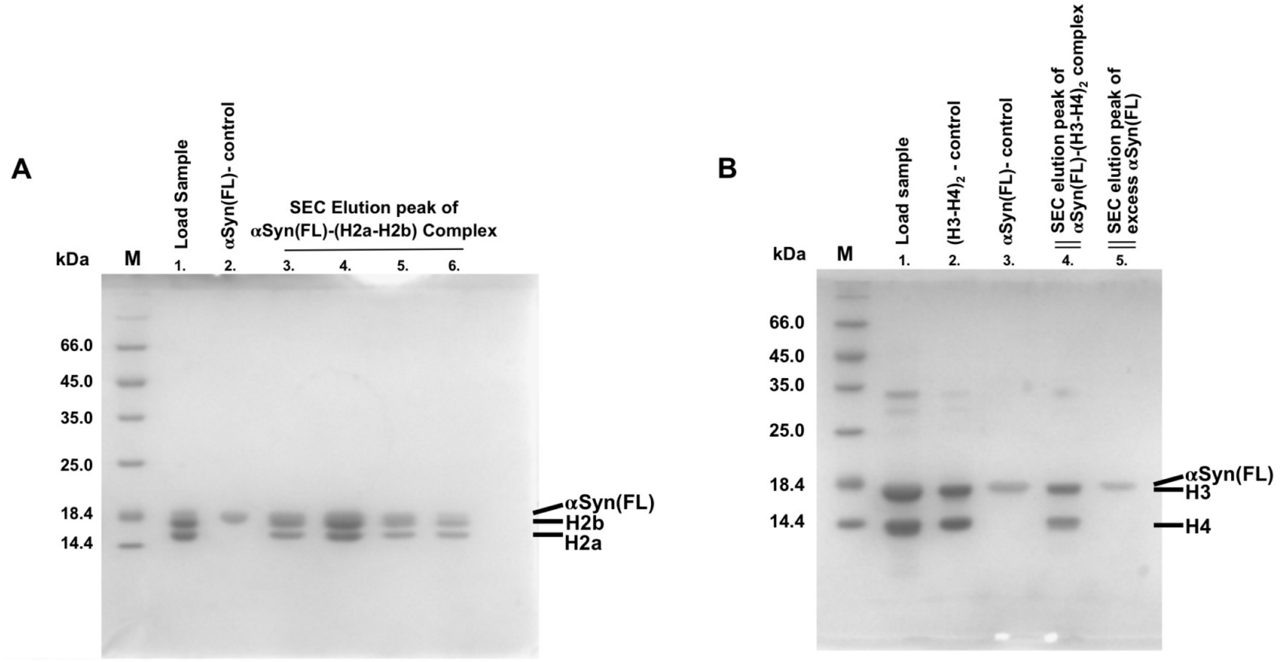


**Supplementary Figure 1:** SDS-PAGE gels indicates the size exclusion chromatography (SEC) elution peaks of **α**Syn(FL) complex with **A.** (H2a-H2b) dimer and **B.** $(H3\text{-}H4)_2$ tetramer. The samples from both SEC were analysed on 15% SDS-PAGE gels. M: Protein ladder.

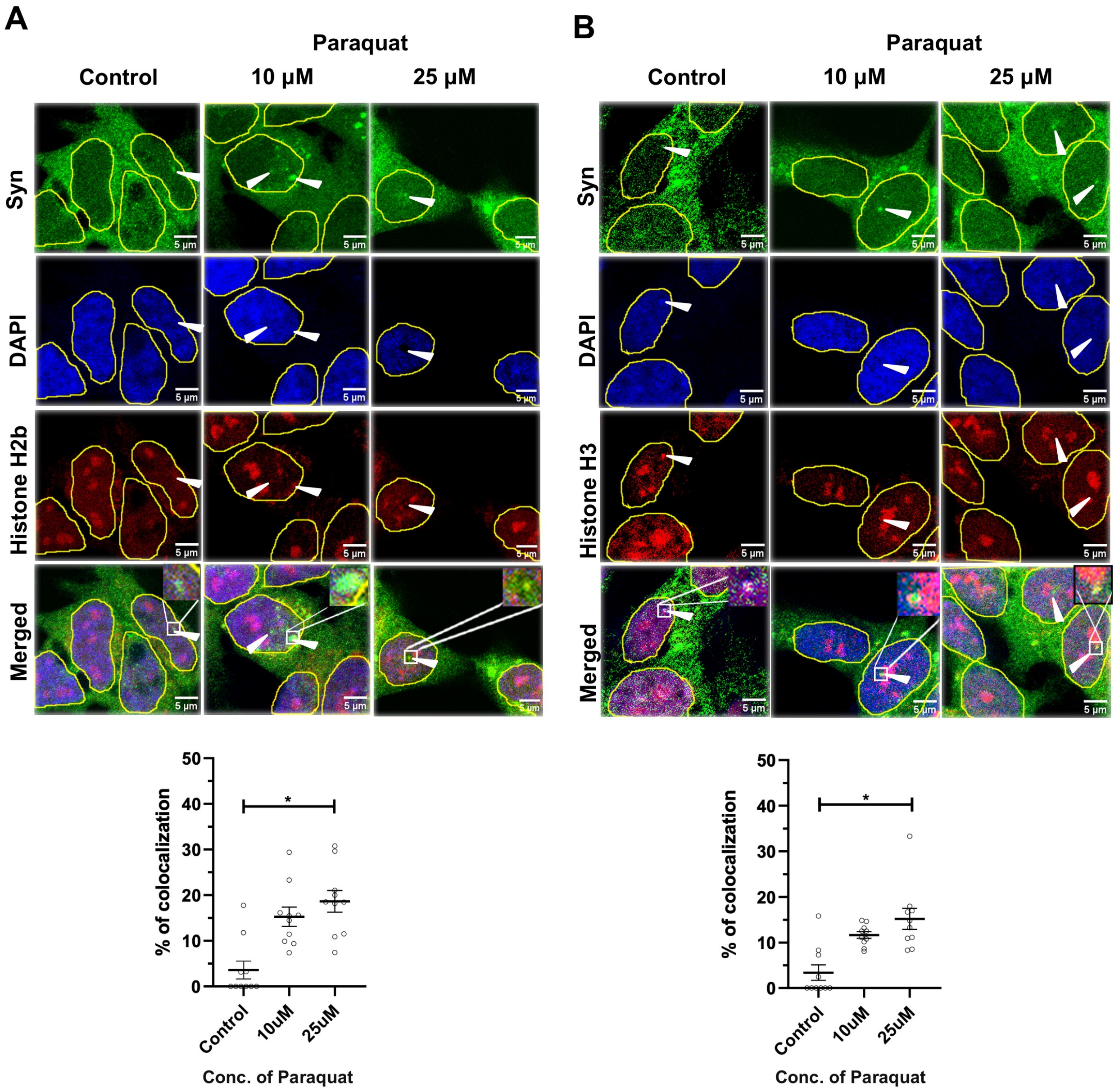


**Supplementary Figure 2: Subcellular localization of SH-SY5Y cells expressing endogenous αSyn imaged using confocal microscopy.** Representative images of endogenous αSyn (green) immunostained against **A.** histone H2b (red) and **B.** histone H3 (red) specific antibodies, DAPI (blue), and merged image. Scale bar: 10 µm. The percentage of histone-αSyn co-localizations for both control and cells treated with 10 and 25 **µ**M concentrations of Paraquat is represented at the bottom ($n = 3$). Quantifications were done from at least 2 images from each independent set, and a total of 10 image frames were analyzed. One-way ANOVA was performed to assess the significant difference between control and treated cells in (A) and (B); $*p < 0.1$.

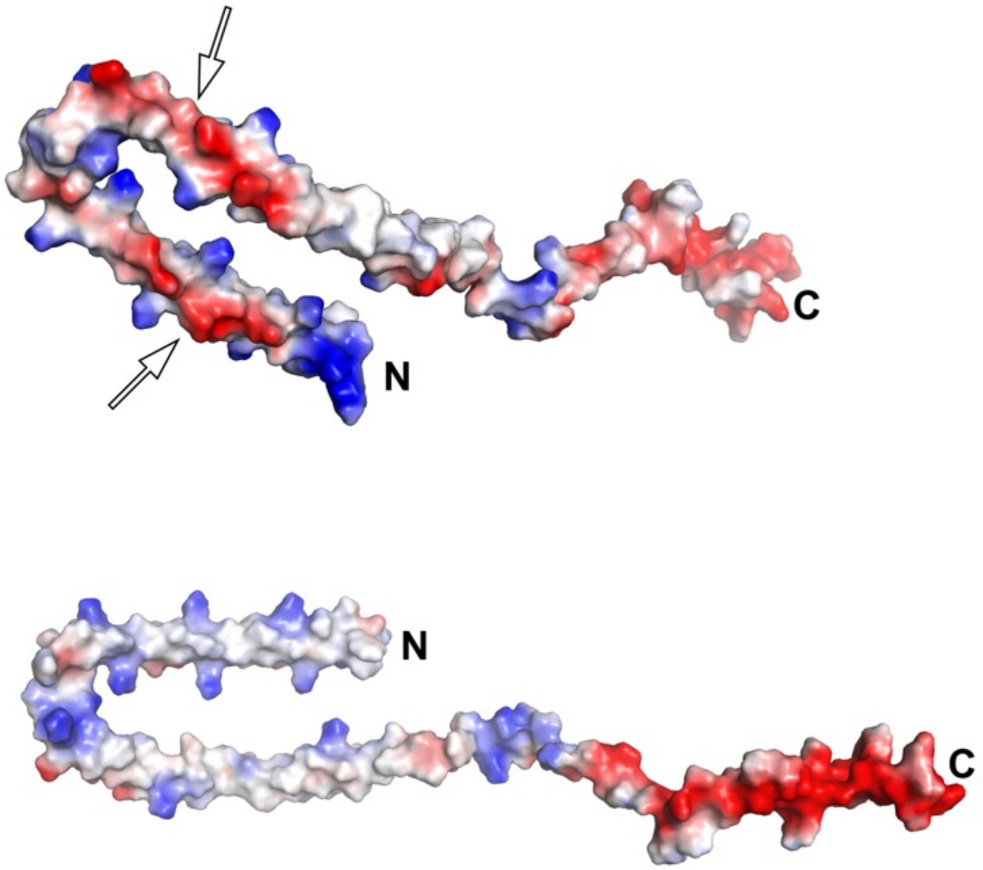


**Supplementary Figure 3: Electrostatic surface representation of αSyn(FL) (PDB: 1XQ8 )** showing amphipathic helices at the N-terminal region (1-103) has negatively charged residues aligned on one side pointed by the arrow (top), and hydrophobic residues on the other (bottom).

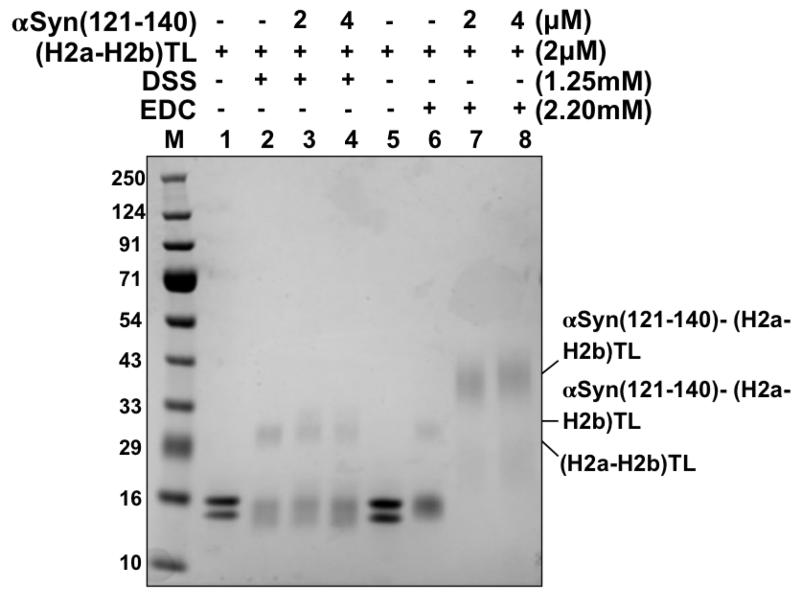


**Supplementary Figure 4:** EDC and DSS mediated cross-linking assays of αSyn(121-140) peptide with (H2a-H2b)TL dimer analyzed on 15% SDS-PAGE gel. Cross-linking studies were performed in the presence of 150 mM NaCl. M: Protein marker along with the indicated protein concentrations at the top.

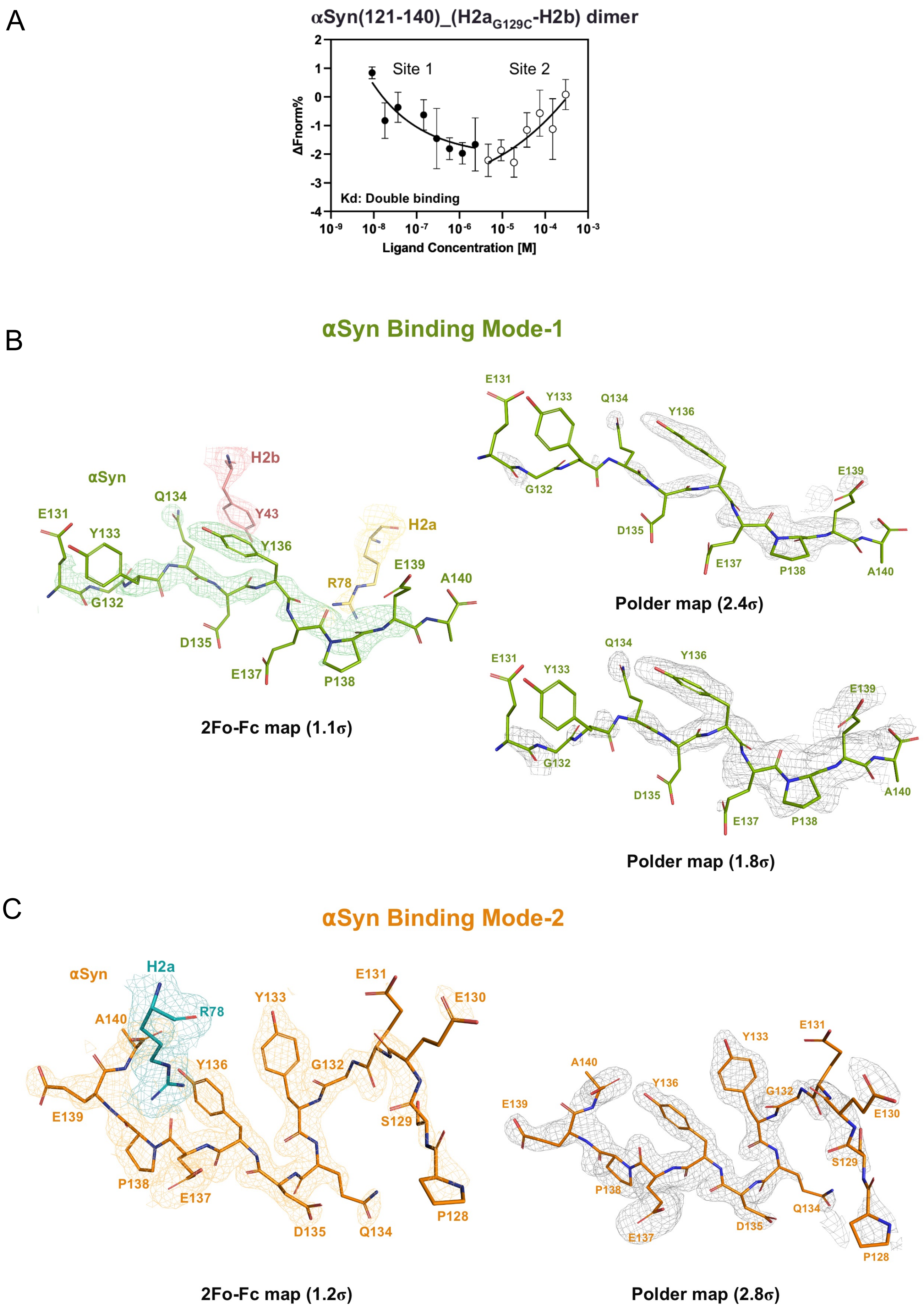


**Supplementary Figure 5: A.** MST analysis of αSyn(121-140) with fluorescently labeled $H2a_{G129C}$-H2b dimer indicates double binding. Error bars represent SD (N=3). **B-C.** 2Fo-Fc and polder maps of **α**Syn binding modes 1 and 2. In binding mode 1, the **α**Syn E131, Y133, D135, and E137 residues side-chain occupancy kept 0.

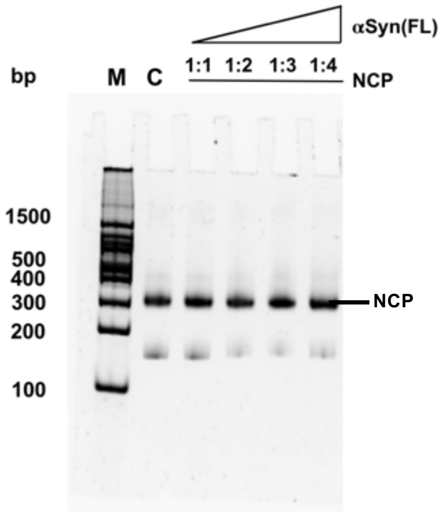


**Supplementary Figure 6:** Electrophoretic mobility shift assay to analyse the interaction between NCP with **α**Syn(FL). The experiment was performed with a fixed concentration of NCP while varying the concentration of **α**Syn(FL). However, no shift in mobility was observed indicating no binding between the two molecules. Control (C) : NCP alone. M: DNA 100bp ladder.
